# The contribution of mate-choice, couple convergence and confounding to assortative mating

**DOI:** 10.1101/2022.04.22.489170

**Authors:** Jennifer Sjaarda, Zoltán Kutalik

## Abstract

Increased phenotypic similarity between partners, termed assortative mating (AM), has been observed for many traits. However, it is currently unclear if these observations are due to mate choice for certain phenotypes, post-mating convergence, or a result of confounding factors such as shared environment or indirect assortment. To dissect these underlying phenomena, we applied Mendelian randomisation (MR) to 51,664 couples in the UK biobank to a panel of 118 phenotypes under AM. We found that 54% (64 of 118) of the tested traits had a causal relationship between partners, with female-to-male effects on average being larger. Forty traits, including systolic blood pressure, basal metabolic rate, weight and height, showed significantly larger phenotypic correlation than MR-estimates, suggesting the presence of confounders. Subsequent analyses revealed household income, overall health rating, education and tobacco smoking as major overall confounders, accounting for 29.8, 14.1, 11.6, and 4.78%, of cross-partner phenotypic correlations, respectively. We detected limited evidence for couple-correlation convergence (e.g. increased similarity with respect to smoking and medication use), measured by stratifying couples by their time spent together. Finally, mediation analysis revealed that the vast majority (>77%) of causal associations between one trait of an individual and a different trait of their partner is indirect. For example, the causal effect of the BMI of an individual on the overall health rating of their partner is entirely acting through the BMI of their partner. In summary, this study revealed many novel causal effects within couples, shedding light on the impact of confounding on couple phenotypic similarity.

## Introduction

In human populations, phenotypic similarity exists between partners compared to random pairs, a phenomenon known as (positive) assortative mating (AM). This has been observed across a wide variety of traits, including anthropometric measures (such as BMI and height), socioeconomic factors, various behavioural and lifestyle measures, (including diet, smoking habits, hobbies, among others), and even disease risk^1–8^. The observed phenotypic similarity can be explained by several factors. First, people tend to and actively seek out partners who are more similar to themselves with respect to certain phenotypes^9,10^. Second, phenotypic similarity can reflect post-mating convergence, where traits become more similar, due to shared household and/or partner influence and interaction over time^11–13^ Finally, non-random assortment with respect to a phenotype can be due to confounders (at the moment of mate choice) such as shared (sociocultural) environment, or geographical barriers^14–16^ Indirect assortment can be viewed as a special case of the latter phenomenon, whereby the confounder is the correlated trait for which direct assortment occurs^17^. As a consequence of 18 assortative mating, the genome of an individual can predict the traits of their partner^18^. Another study even found evidence of direct genetic associations between the genome of an individual and the traits of the partner, suggesting that partner heritability of a trait cannot be solely explained by between partner trait correlation^19^. The causes and consequences of phenotypic assortment remain unresolved and have implications in the study of human behaviour, population genetics, and public health. For instance, increased phenotypic similarity could naturally imply genetic similarity, leading to variants that are otherwise independent to become correlated, and ultimately resulting in a concentration of (genetic) resources^20–22^.

Any trait influenced by shared confounders will show assortment. Therefore, it is crucial to separate traits under direct assortment from those that show partner-similarity due to being driven by another trait/factor under direct assortment (Figure 1). Thus, we identify three underlying phenomena leading to AM: (i) traits can be under direct- and/or (ii) indirect mate choice, and (iii) additionally modified by post-mating convergence (during cohabitation). These phenomena can be rephrased for modelling purposes as follows: direct assortment is treated as a (direct) causal effect acting between the traits of the couple (index to partner) and indirect assortment as a confounder effect. Confounder effects can emerge due to shared factors (e.g. socio-economic, geographic) and/or driven by correlated traits under direct assortment, and these can occur pre-mate choice (or intensified post-mate choice due to shared household/habits).

**Figure 1:**
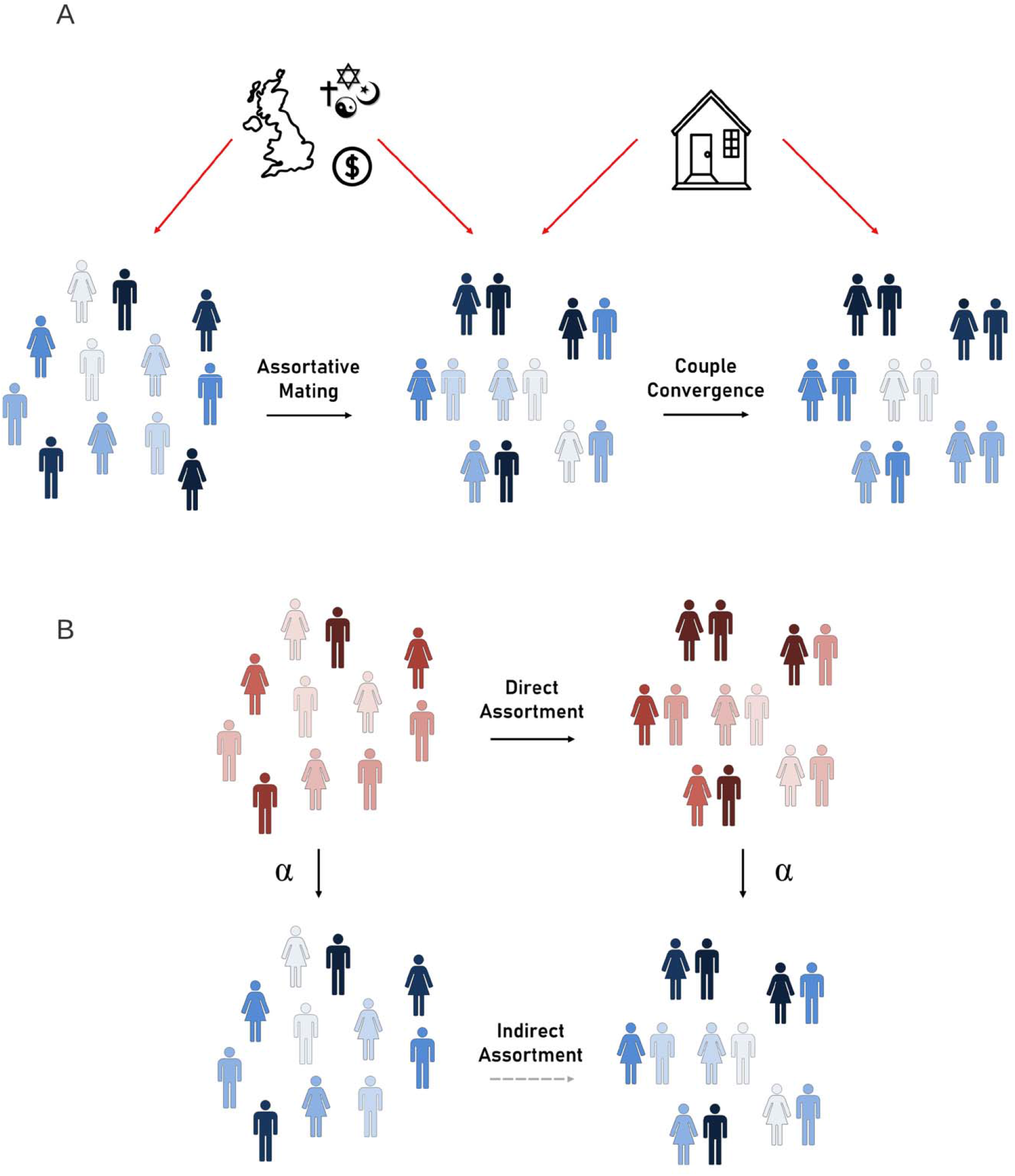
Assortative mating framework. **A.** Illustrates a trait (given by the colour blue) which is under assortative mating, either directly (through mate choice) or due to confounding factors such as shared geography, cultural or religious status or socioeconomic measures. Subsequently, this trait may also undergo post-mating convergence which could be due to direct causal influence from one partner on the other (i.e. through imitation or influence) or due to confounding factors such as shared environment. **B.** Illustrates a trait which shows increased similarity among couples (given by the blue trait), however this assortment is only observed because of a causal effect that exists between another trait (shown in red) acting on the blue trait. For example, if direct assortment occurs under a trait such as BMI (i.e. couples intentionally select partners of similar BMI as themselves), phenotypic correlation will also be observed at all traits which have a causal effect of BMI, such as blood pressure, fasting glucose, etc.

Despite some pioneering work, it remains difficult to untangle the role of the three outlined components in explaining observed phenotypic similarity between partners and resolving the impact of confounding from casual factors. Analogous to classical epidemiological studies, where it is difficult, if not impossible, to discern causal factors from confounders, mere phenotypic similarity among couples is susceptible to the same interpretational limitations and challenges. Mendelian randomization (MR) is an alternative approach which is used to assess causality leveraging large-scale observational data (including genetics). MR takes advantage of the random allocation of genetic variants at birth to infer causality between an exposure and an outcome^23^, thereby minimizing the possibility of reverse causality and confounding. To date, MR has proven to be a reliable causal inference method, revealing thousands of novel, causal relationships between exposures and outcomes.

In this work, we sought to adapt MR by examining causality between individuals, where the exposure and outcomes traits are measured in different individuals (whereas classical MR designs involve a single individual, e.g. BMI risk on CAD). A similar approach has been attempted for exploring couple effects with respect to alcohol consumption, and it was shown that while the observed phenotypic correlation in couples does not tend to increase with age, the couple correlation and the estimated direct causal effect differed substantially^24^. Here, we examined a large number of complex traits and applied MR to estimate the direct causal effects impacting matechoice, explored the impact of time couples have lived together on their phenotypic similarity, and examined the cumulative role of a wide range of potential confounders on trait correlations between partners. Finally, we explored how cross-trait AM emerges by dissecting them to direct and indirect (same-trait AM combined with classical (same-sample cross-trait) causal effects) counterparts.

## Methods

### Sample selection and couple definition

This study used the UK Biobank (UKBB) cohort, a prospective population-based study with over 500,000 adult participants. Couples were identified and selected according to the following procedure. The initial UKBB sample comprised 502,616 individuals. First, participants were filtered to only genotyped, white, unrelated individuals according to the genetic QC file (specifically participants were retained if they had the following values in the QC file: “excess.relatives” = 0, “putative.sex.chromosome.aneuploidy” = 0, “in.white.british.ancestry.subset” = 1 and “used.in.pca.calculation” = 1). Redacted samples and participants that removed consent were also excluded. After filtering, 337,138 participants remained. Within this sample, we retained individuals coming from households with exactly two unrelated, opposite-sex individuals, leaving 108, 898 participants. Finally, using the data at data-field 6141, “How are people in household related to participant” pairs were filtered to only include couples who had both responded “Husband, wife, or partner”, leaving 103,328 participants, or 51,664 couples for downstream analyses (Supplementary Figure 1).

### Mendelian Randomisation

MR uses genetic variants as instrumental variables (IVs) to assess the presence of a causal relationship. The random distribution of genetic variants at birth reduces the possibility of confounding or reverse causation as explanations for the link between the exposure and outcome in the same way that the random allocation of a therapy in a randomized controlled trial minimizes this risk. MR relies on three core assumptions for the genetic variants. First, IVs must be associated with the exposure of interest (the relevance assumption). Second, IVs must not be associated with any confounder in the exposure-outcome relationship (the exchangeability assumption). Third, IVs must not affect the outcome except through the exposure (the exclusion restriction assumption). There are several methods to estimate the causal effect using MR, the simplest being the Wald method, whereby a ratio is taken between the variant-outcome association and the variant-risk factor association. A natural extension of this approach, known as the inverse-variance weighted (IVW) method, combines multiple IVs, applied in this report^25^. The causal effect of exposure *X* on the outcome *Y*, using *k* genetic variants, is given by

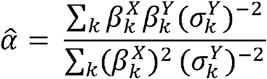

with the corresponding variance 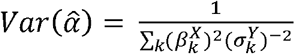, where 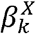 and 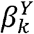 represent the estimated effects of genetic variant *k* on *X* and *Y*, respectively and 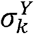 represents the standard error of 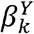.

### Phenotype selection and processing

We used an agnostic, phenome-wide approach for selecting phenotypes. Specifically, we first selected phenotypes which were analysed by the Neale group and which had both male, female and joint summary statistics available (http://www.nealelab.is/uk-biobank/). This list was intersected with our internal database (application number #16389), leaving 1,278 phenotypes available for analysis. Phenotypes were processed in the filtered QC-data set (N = 337,138) according to a slightly modified version of the PHESANT pipeline to accommodate the phenotypes that we had available in our database^26^. Continuous variables were transformed to a normal distribution using a rank-preserving inverse normal quantile transformation (INQT), while ordinal and binary traits were re-categorized according to PHESANT documentation (for e.g. categories with less than 10 participants were removed). We then filtered these phenotypes as follows. First, in order to focus on traits with some indication of assortment, we computed the raw phenotypic correlation amongst couples and removed phenotypes with a Pearson correlation < 0.1. To ensure that INQT was not significantly impacting the correlations of each trait, we also calculated the correlation between partners for each trait using the non-parametric Spearman correlation and found consistent estimates (Supplementary Figure 2). Second, we removed phenotypes which had less than 5 valid IVs for MR. IVs were defined based on an association *p* < 5 × 10^-8^ in the joint Neale summary statistics, after pruning for independence (based on a clumping procedure performed in PLINK with the options --clump-kb 10000 and --clump-r2 0.001 using the 1000 Genomes European samples as a reference). Third, using the sex-specific summary statistics, the IV heterogeneity between sexes was calculated. IVs that showed (Bonferroni corrected) significant evidence of heterogeneity between sexes were excluded (*p* < 0.05/[number of IVs]). After this procedure, phenotypes were again filtered to those with at least five valid IVs remaining. Fourth, dietary phenotypes were removed due to high correlation amongst these phenotypes (due to the shared household), insufficient power, problems with reverse causation and difficult interpretation^27^. Finally, we manually removed several duplicated and redundant phenotypes. Specifically, (i) left-side body traits (highly correlated with right-side) were removed, (ii) we retained only one of the duplicated phenotypes for BMI and weight (retaining UKBB data fields 21001 and 21002, respectively), and (iii) all “qualifications” data was removed (corresponding to UKBB field <u>6138</u>) due to the availability of finer-scale correlated variables, such as “age completed full time education” (data field 845). After this process, 118 phenotypes remained for analysis (see Supplementary Figure 3).

### Estimation of single-trait causal effects in couples

To investigate the causal effect of a trait in one individual on the same trait of their partner, we performed couple-specific MR analyses. Specifically, the trait in the index case was used as the exposure, and the same trait in the partner was used as the outcome trait. The effect of genetic variants on the exposure were obtained from the Neale summary statistics, using the full UK Biobank sample. Instruments for each trait were selected as described above, i.e. being both genome-wide (GW) significant (*p* < 5×10^-8^) and pruned for independence. Next, we estimated the effects of SNPs on the outcomes of interest by testing the association between each genetic instrument measured in the index individual with the phenotype measured in the partner using the UKBB partner data set described above. In other words, for each phenotype, the corresponding genetic data for the IVs were obtained from the index case while the phenotypes (dependent variable) were taken from the corresponding partner. All SNP-trait estimates were estimated in males and females separately (i.e. using the sex-specific Neale summary statistics or two separate models in the couple data), adjusting for age and the first 40 genetic principal components (PCs) of both the index and partner. To mimic the Neale models, we performed linear regression of SNP effects on phenotypes, regardless of data type (including binary). Continuous phenotypes were scaled to have mean 0 and SD of 1 before regression, while ordinal and binary phenotypes were left as processed by PHESANT.

To estimate the causal effect of a trait from an index case to a partner (*α*_*x_i_*→*x_p_*_), we combined the effects of genetic instruments on the exposure (from Neale) with effects on the outcomes (measured among couples) in an MR framework using the IVW method (Figure 2A)^25^. To estimate the causal effects in both sexes combined, SNP-effects were first meta-analysed across sexes using fixed effects models prior to performing MR (rather than meta-analysing the MR estimates directly) to minimize weak instrument bias^28^. Effects of the genetic estimates on both the exposure and outcome were first standardized (such that the squared effect size represents the explained variance) to allow for seamless comparison across traits and to the raw phenotype correlation. Significance was determined by adjusting for the number of effective tests based on the correlation matrix of phenotypes tested^29^, resulting in 66 independent tests. The significance threshold was adapted accordingly as *p* < 0.05/66.

**Figure 2:**
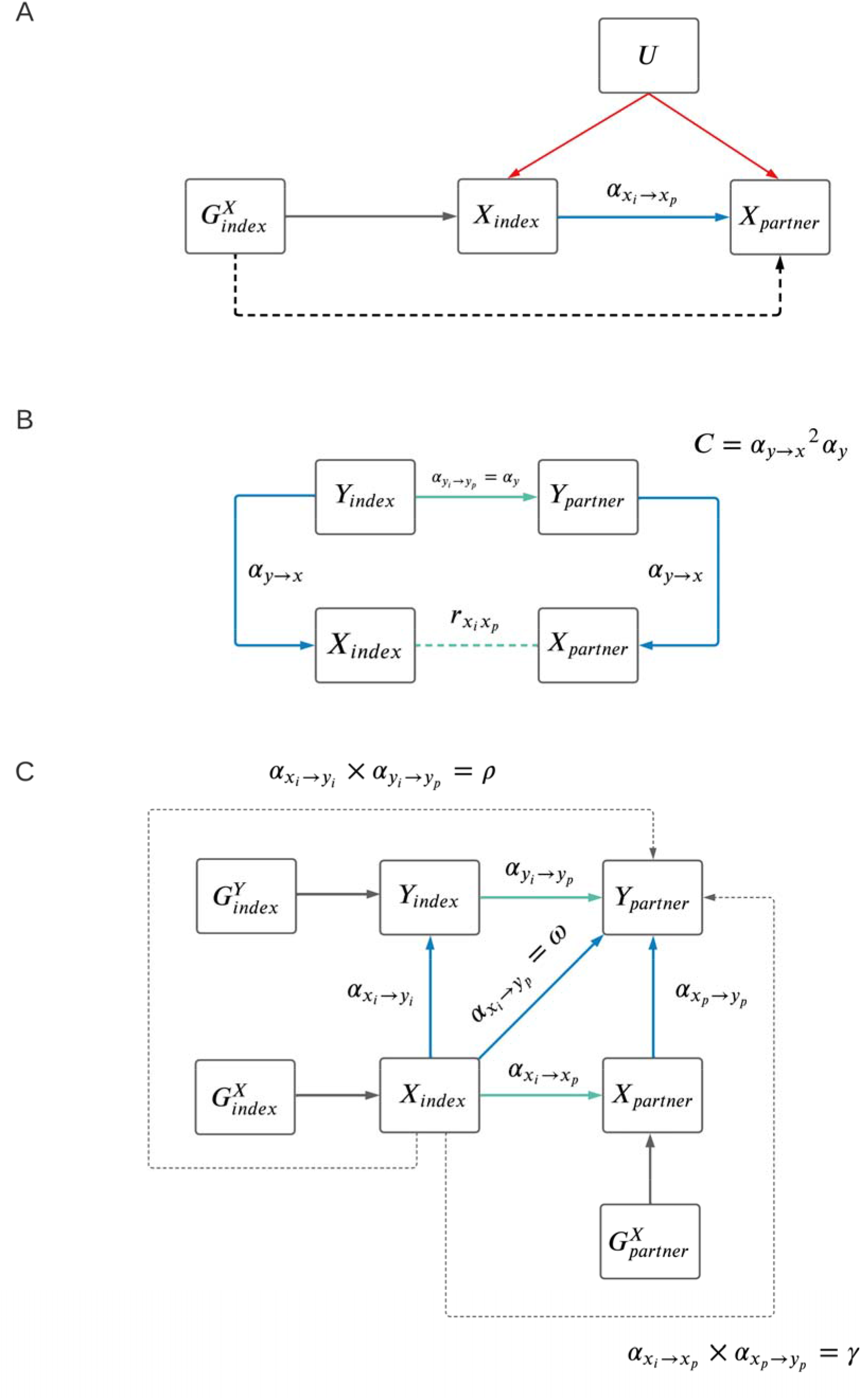
Mendelian randomisation schematic within couples. **A.** Illustrates the causal effect among couples with a single trait (α_x_i_→x_p__), where G represents genetic variant(s), X represents a single trait (in an index and a partner), and U represents confounding factors which are not associated with genetic variance owning to the random distribution of alleles at conception. **B.** DAG illustrates the impact a confounder (trait Y) could have on the phenotypic correlation between partners for a given trait X (r_x_i_x_p__). Correlation due to confounding can be calculated as C = α_y→x_^2^ × α_y_. **C.** Represents the expanded causal network involving two traits and the various estimated causal paths from trait X of an index case (X_i_) to a phenotype Y in the partner (Y_p_) given by ω, γ, and ρ. Cross-trait causal effects from X_i_ to Y_p_ (ω) can be summarized by three possible (non-independent) scenarios: (1) X_i_ could exert a causal effect on X_p_, followed by X_p_ having a causal effect on Y_p_ in the partner alone (γ); (2) the reverse could occur whereby X_i_ has a causal effect on Y_i_ in the index alone, followed by a causal effect of Y_i_ case on Y_p_ (ρ); (3) there could be other mechanisms, either acting directly or through other unmeasured/considered variables. To quantify p, we first estimated the causal effect of Y_i_ on Y_p_ in multivariable MR (not illustrated), to exclude any residual effect of X on phenotype from index to partner. These three scenarios could also act in some combination. Therefore, the ω estimate would capture the paths of γ, ρ and other mechanisms combined. In both **A** and **C**, cross partner causal effects are given by blue arrows, and same-person causal effects are given by green arrows.

After estimating single trait causal effects in couples, we used a two-tailed z-test to identify traits with a significant difference between the MR-estimate and the phenotypic correlation in couples. For each of trait with discrepant estimates, we tested the causal effect of each of the remaining phenotypes in our pipeline (*Y*_1_,…, *Y*_117_) on the focal trait of interest (*X*) using MR (*α*_*y*→*x*_). These same-person MR estimates were calculated using meta-analysed sex-specific Neale estimates for both the SNP-exposure and SNP-outcome effects using the IVW-method. Before performing each same-person MR, genetic variants were first filtered for evidence of reverse causality at a threshold of *p* < 0.001 (Steiger filter)^30^, whereby SNPs were removed if the standardized SNP effect on the outcome was stronger than the effect on the exposure based on a one-tailed t-test at a significance level of p < 0.001. SNP-effects were standardized prior to calculating MR effects.

We then explored those potential confounders, *Y_k_*, with a significant impact on *X* (*p* < 0.05/66). As the confounding impact of each *Y_k_* involves a within couple effect (*a*_*y_i_*→*y_p_*_), as illustrated in Figure 2B we further filtered the remaining *Y_k_* traits, to those with a significant within couple MR effect (*p* < 0.05/[number of remaining *y_k_*]). After identifying potential confounder traits, we combined these *α*_*z*→*x*_ with the within couple causal effect (*α*_*y_i_*→*y_p_*_), and calculated the correlation due to confounding as *C* = *α*_*y*→*x*_^2^ × *α*_*y_i_*→*y_p_*_ to determine the contribution each trait (*Z*) confounds the within couple correlation for trait *X* (*r_x_i_x_p__*). We subsequently calculated the ratio of this correlation (*C*) and the correlation of *X* in partners as *C*/*r_x_i_x_p__*.

### Assessing the role of confounders on trait correlation in couples

We sought to explore the impact of potential confounders on mate-choice by calculating the trait correlations between partners that are due to confounding. We considered the impact of the following confounders (*Y*) on the partner correlations of the remaining 117 traits selected by our pipeline: average household income, age completed full-time education, sports club or gym user, current tobacco smoking, overall health rating and North and East birth place coordinates (UKBB data fields 738, 845, 6160, 1239, 2178, 129, and 130 respectively). Using the single-trait causal effects in couples and the same-person MR-estimates, correlation due to founding was calculated for each pair (*Y, X*) as *C* = *α*_*y*→*x*_^2^ × *α*_*y_i_*→*y_p_*_ (Figure 2B). These confounding estimates were finally contrasted to the couple correlation values to explore the extent that each Z may confound couple correlations by examining the ratio between the two estimates (i.e. *C*/*cor*(*X_i_, X_p_*)). Birthplace coordinates (east-west and north-south) were considered together, and their invoked trait correlations were summed up, as they are orthogonal by definition.

### Investigating the effect of time and age on correlations and causal relationships in couples

Trait similarity in couples can be driven by both mate choice and/or trait convergence over time spent together. To tease out the contribution of these different sources, we explored whether the cross-partner causal effects change as a function of the length of the relationship and age. The length of relationship was proxied by the minimum value of the “length of time at current address” (data field 699) for the two partners. To estimate the effect of age, we took the median age of couples. For each of the two derived variables, we split the couples into five roughly equal sized bins (using the “smart_cut” function from the cutr R package). We first estimated the phenotypic correlation of each trait, within couples of each bin. Next, for each single-trait MR described above, analyses were run in the full sample as well as in the different bins. Of the significant results identified in the sex-combined analysis above, we tested to see if there was any significant difference in MR-estimates amongst the bins. Binned MR-estimates were computed using SNP-outcome effect estimated in each bin separately, and the SNP-outcome effects used the same SNP-exposure effects from Neale. Analyses were run in each sex separately and combined (meta-analysed at the SNP level). As above, SNP effects were standardized prior to calculating MR estimates. To assess for the presence of a trend across bins, we tested the significance of the slope of a linear model of bin-specific correlations and bin-specific MR-estimates, inversely weighted by the SE, versus the bin centre (i.e. the median age or time-spent-together for the given bin). Multiple testing was, as described above, adapted based on the effective number of tests, but restricted to traits which showed significant causal effects in the joint (both sexes combined), non-binned MR (resulting in a threshold of *p* < 0.05/29).

### Estimation of cross-trait causal effects in couples

Using the same process as in the AM analysis involving a single trait, we also sought to investigate causal effects within couples involving two traits (*α*_*x_i_*→*y_p_*_). In other words, two different traits were used as exposure and outcome to determine the causal effects of trait *X* (in the index individual) on trait *Y* (in the partner). Here, we only considered trait combinations with phenotypic correlation < 0.8 (estimated in the entire UKBB, N = 337,138), in order to avoid too closely related traits. The same set of SNPs were used as in the same-person MR (i.e. first filtered for the presence of reverse causality). As in the single trait MR, SNP-exposure effects were obtained from the Neale summary statistics and SNP-outcome effects were estimated in the couple derived dataset. MR models were run in both sexes separately and jointly (meta-analysing the SNP effects before performing MR analyses). Significance was determined based on the squared effective number of tests (*p* < 0.05/[66^2^]).

### Comparison of paths from index to partner

There are several independent paths through which a trait in an index case could exert a causal effect on another trait in the partner. We wanted to explore if one path was more dominant, in general, and if there was evidence for the presence of other (confounder) traits involved. Restricting to Bonferroni-significant trait pairs (with phenotypic correlation < 0.8) from the couple MR, we sought to explore the various paths through which a phenotype *X* in an index case (*X_i_*) could causally impact a phenotype in the partner (*Y_p_*) as illustrated in Figure 2C. With the exception of exposure traits that directly alter the environment of their partner, such as smoking creating the presence of second-hand smoke, *X_i_* is unlikely to have a direct effect on another *Y_p_*. Alternatively, *X_i_* might indirectly impact *Y_p_* by inducing changes in *X_p_*, which in turn impacts *Y_p_*. For instance, increased BMI in an index case is not expected to directly increase cardiovascular disease risk in their partner, but rather to modify the partner’s risk through first increasing their BMI. To explore this intuition, we dissected the causal effect from *X_i_* to *Y_p_* (*ω*) into three possible (non-independent) mechanisms. First, *X_i_* could exert a causal effect on *X_p_*, followed by *X_p_* having a causal effect on *Y_p_* in the partner alone (y). Second, the reverse could occur whereby *X_i_* has a causal effect on in the index alone, followed by a causal effect of *Y_i_* case on *Y_p_* (*ρ*). Third, there could be other mechanisms, either acting directly or through other unmeasured/considered variables. These three scenarios could also act in some combination. In this way, the *ω* estimate would capture the paths of *γ, ρ* and other mechanisms combined.

Using the same-person MR estimates (*α*_*x*→*y*_) that were calculated as described above, we estimated *γ* and *ρ* representing the various paths from *X_i_* to *Y_p_*. To quantify *γ*, the single-trait couple causal effect estimate (i.e. from the regression *X_p_* ~ *X_i_*) were multiplied by the same-individual causal estimate (i.e. *α*_*x*→*y*_ from *Y* ~ *X*). To quantify *ρ*, we first estimated the causal effect of *Y_i_* on *Y_p_* in multivariable MR (MVMR), to exclude any residual effect of *X* on phenotype *Y* from index to partner. Specifically, *Y_p_* was used as the independent variable with both *Y_i_* and *X_i_* as independent variables (i.e. the MVMR was *Y_p_* ~ *Y_i_* + *X_i_*). We included both IVs from *X* and *Y*, pruned for independence (performed in PLINK with the options --clump-kb 10000 and --clump-r2 0.001 using the 1000 Genomes European samples as a reference). We took the coefficient of *Y_i_* as the direct causal effect from *Y_i_* to *Y_p_* (*α*_*y_i_*→*y_p_*_) and multiplied this by the same-individual causal estimate (*α*_*x*→*y*_). Finally, we estimated *ω* directly from our cross-trait couple MR framework (*α*_*x_i_*→*y_p_*_). We compared the estimates of *γ, ρ*, and *ω* using a z-test to assess their difference and using linear regression with the intercept forced through the origin to determine their relationship. Finally, we quantified the proportion of *ω* that could not be explained by the paths quantified by *γ* and *ρ*. As *γ* and *ρ* are not perfectly independent, potentially due to correlation between *X* and *Y* or pleiotropic limitations of MR, we estimated the extent of dependence via the correlation between *p* and *γ* across the different trait pairs. To account for the duplicate signals due to this correlation, we removed the effects of *γ* from *p* by keeping the residuals from the linear regression p ~ γ. We then estimated the proportion of variance explained (*R*^2^) of *ω* jointly by *γ* and the residualised *ρ*.

## Results

### Effect of sex, age and time together causal effects in couples

Among the 118 phenotypes tested, we identified 64 significant causal effects in partners after adjusting for the effective number of tests (*p* < 0.05/66) (see Supplementary Table 1). We also examined the Cochran’s heterogeneity Q-statistic to identify traits with high heterogeneity and found no evidence of heterogeneity in the MR-estimates (all *p* > 0.05/66). We assessed the 64 significant results for significant sex-differences, but did not identify any after adjusting for the effective number of tests among the remaining traits based on their pair-wise correlation matrix (*p* < 0.05/29). However, 15 traits showed a nominally significant difference between sexes (*p* < 0.05, Supplementary Table 2), which is 4.7-times higher than expected (*p_binomial_* = 7.45×10^-8^). Applying a paired t-test among these 15 traits revealed that female-to-male MR-estimates are on average larger than male-to-female estimates (*p* = 0.014).

To identify if partner traits converge over time, we explored the impact of age and time-spent-together (proxied by the amount of time at same address) among the 64 significant traits in both males and females separately and both sexes combined. Using linear regression of MR-estimates versus the median of the five age and/or time-spent together bins, we detected no significant results in the sex-combined results after adjustment for number of effective tests (p < 0.05/29). We also examined the Pearson phenotypic correlation within the different bins and assessed for the presence of a trend, using linear models (phenotypic correlation versus median bin). Two traits which showed a significant (p < 0.05/29) trend across the bins according to time-spent-together, namely body fat percentage and hand grip strength (right). In both cases, the correlation decreased as time-spent-together increased. We found another two traits which showed a significant trend across the bins by median age, namely smoking status: previous and aspirin use. In this case, for both phenotypes, the slope increased as age median-age increased (Figure 3 and Supplementary Table 3). Consistent results were found using Spearman correlation (all *p* < 0.05/29).

**Figure 3:**
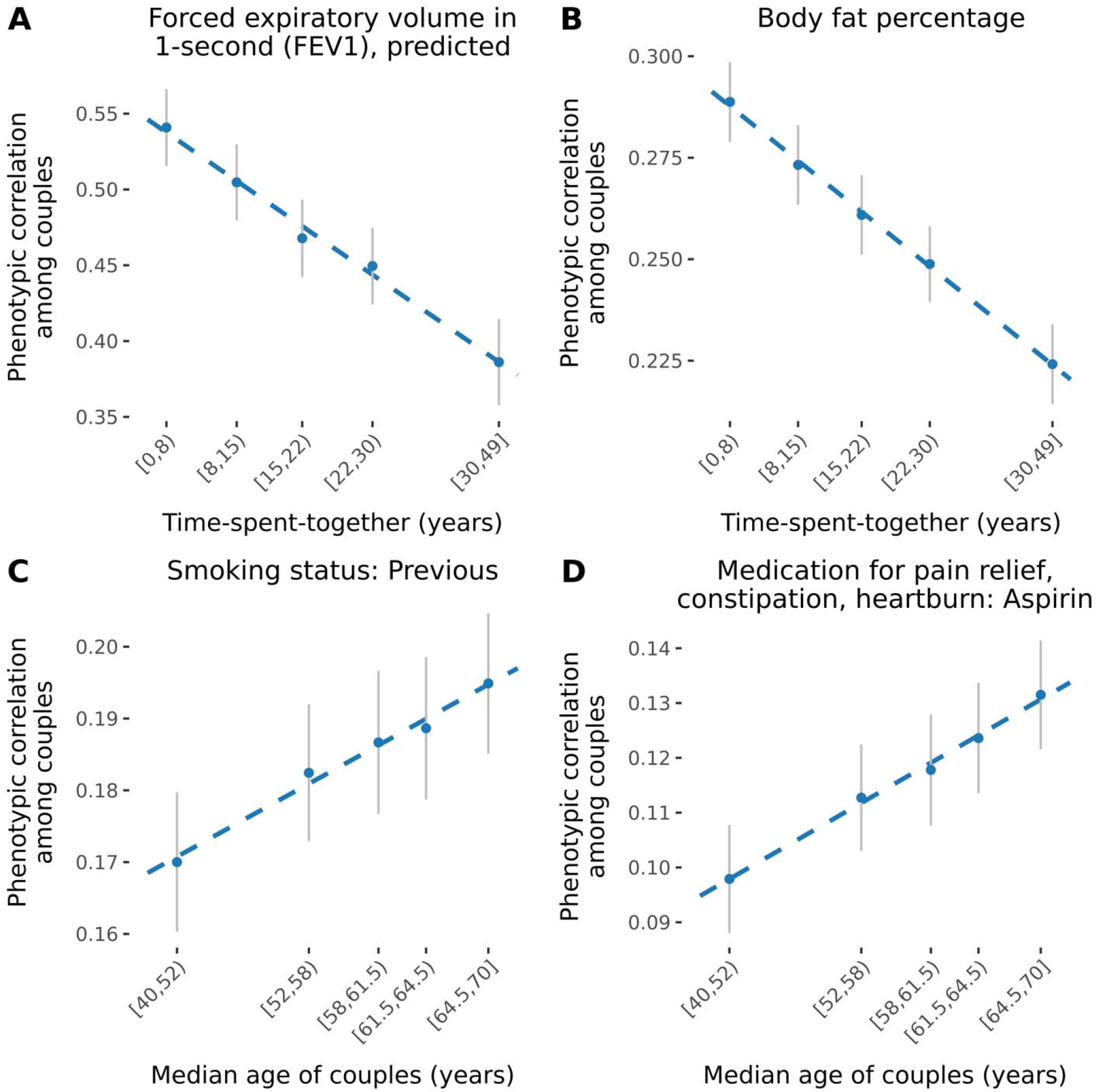
Phenotypic correlation for selected traits by time-spent-together and age of couples. Scatter plots show the phenotypic correlation among couples within different bins. Couples were binned by time-spent-together (proxied by the time lived at same household, panels **A** and **B**) and median age (panels **C** and **D**).

### Relationship between causal effects and raw phenotypic correlation in couples

To better understand the nature of phenotypic assortment, we assessed whether there were any discrepancies between the causal effects within couples and observational correlations. Using MR, estimates for the causal effect from index to partner within couples were estimated for 118 phenotypes, selected based on their elevated correlation between partners and sufficient [> 5] valid IVs rendering them suitable for MR analysis. Using a two-tailed Z-test to gauge the statistical significance of the difference between the estimates, we compared (standardised) causal MR effects to the raw phenotypic correlation among couples to identify any traits where the correlation was different than the MR-estimate. After adjusting for the effective number of tested traits (p < 0.05/66), we identified 43 traits which showed different phenotypic correlation compared to MR-estimate (see Figure 4A, Supplementary Table 4). Of these, three had a larger MR-estimate compared to correlation (time spent watching television, comparative height size at age 10, and overall health rating), while the remaining 40 traits had a larger correlation compared to MR-estimate. Among these included place of birth, North-coordinate (NC) (*r* = 0.58 *vs α* = 0.33); systolic blood pressure (SBP) (*r* = 0.16 *vs α* = 0.05); height (*r* = 0.25 *vs α* = 0.21); forced vital capacity (FVC) (*r* = 0.25 *vs α* = 0.13); basal metabolic rate (BMR) (*r* = 0.21 *vs α* = 0.16); and basophil count (*r* = 0.47 *vs α* = 0) (see Figure 3).

**Figure 4:**
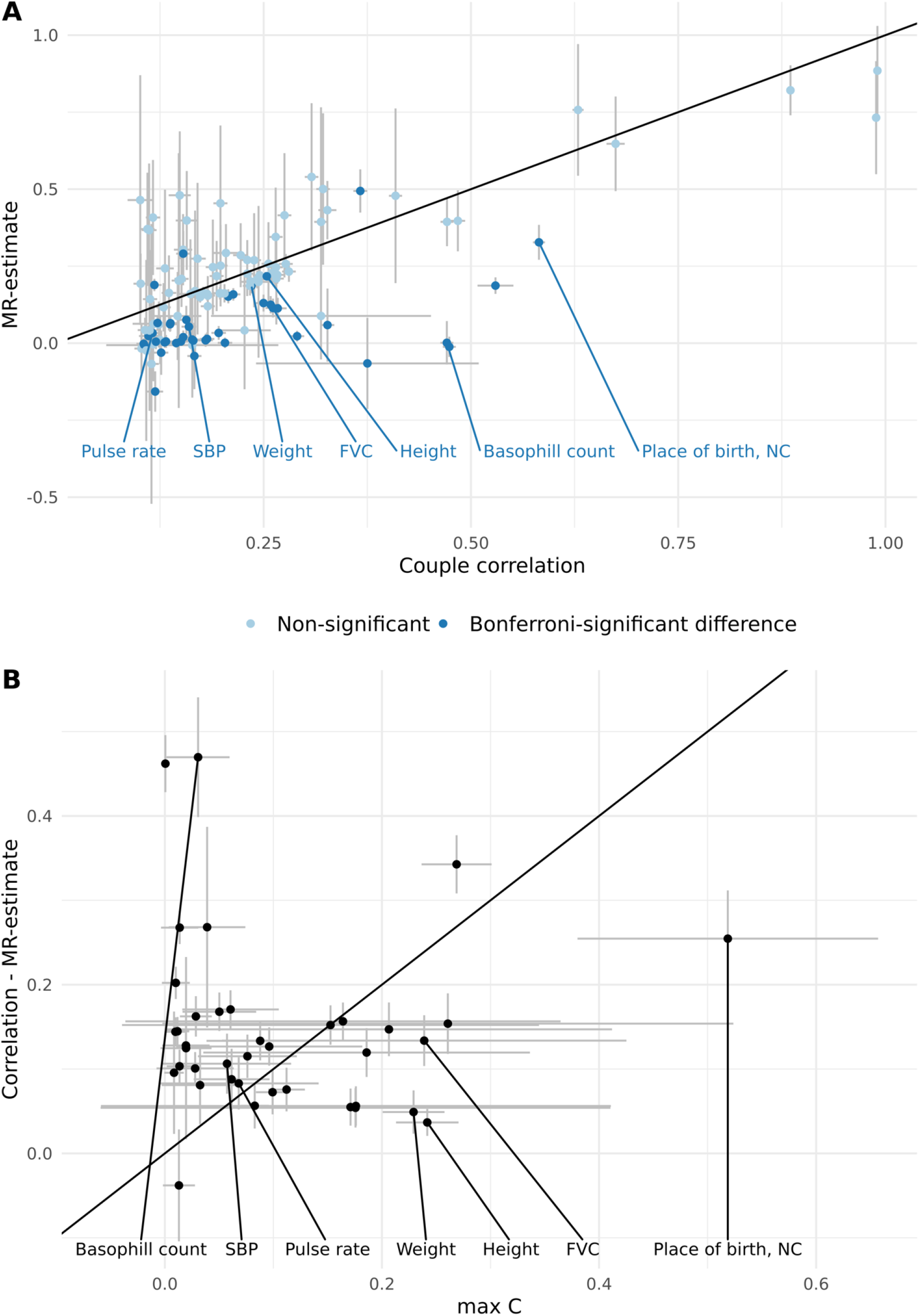
Phenotypic correlation in couples versus causal effects and evidence of confounder traits impacting the discrepant estimates. **A.** Scatter plot shows the within couple standardized MR-estimates (*α*_*x_i_*→*y_p_*_) versus the phenotypic correlation among couples (*r_x_i_x_p__*); error bars represent 95% CIs. A two-tailed z-test was used to test for a significant difference between the estimates. After adjusting for the number of effective tests (p < 0.05/66), 43 significant differences were identified (shown in dark blue), where 3 traits showed larger MR-estimates compared to correlation, and 40 traits showed larger correlation compared to MR-estimates. The identity line is shown in black. Labelled pairs are discussed in the main text. Abbreviations: SBP: systolic blood pressure; FVC: forced vital capacity; NC: North coordinate. **B.** Scatter plot shows the difference in phenotypic correlation and MR-estimate versus the maximum C for each trait where the phenotypic correlation was greater than the MR-estimate (number of traits = 39); error bars represent 95% CIs. The identity line is shown in black. Abbreviations: SBP: systolic blood pressure; FVC: forced vital capacity; NC: North coordinate.

Significant differences could be indicative of the presence of confounders (either negative or positive) driving the observed phenotypic correlation. Thus, for traits where couple correlation was significantly different than MR causal estimates, we sought to identify potential confounders which may, in part, explain the discrepant estimates. For the three traits where correlation was less than MR-estimate, we searched for negative confounders (i.e. negative *α*_*y_i_*→*y_p_*_), but did not identify any. Conversely for traits where the correlation was greater than the MR-estimate we searched for positive confounders, and found many potential positive confounders. Namely, the mean number of potential confounders from our set of 117 candidates was 22.56, with a maximum of 39, for only one trait we did not identify any potential confounders (Supplementary Table 5). For instance, for systolic blood pressure, we identified 29 (correlated) potential confounders which may explain the larger phenotypic correlation as compared to MR effect. These potential confounders included physical activity, BMI, lung fitness measures, overall health rating. For weight, we found 30 potential confounders, including anthropometric traits (such as leg, trunk, arm fat mass), various behavioural traits which are reflective of exercise patterns, such as time spent watching television, walking pace, phone use, among many others (Supplementary Table 5). Many of the 40 traits with larger phenotypic correlation compared to MR-estimates included blood cell counts and/or percentages (such as white blood cell (leukocyte) count, neutrophil count, monocyte count and percentage, reticulocyte percentage and count). The potential confounders for these traits were highly overlapping, including physical activity level, anthropometric traits, smoking and health rating (Supplementary Table 5). Other notable confounders included measures of physical activity for forced vital capacity; smoking status and fitness measures for basal metabolic rate; and measures of body size for hand grip strength. Finally, for each confounder we calculated the correlation due to confounding (C) as described above (see Figure 1B). We then compared the difference in estimates to the maximum C for each trait (Figure 4B) since the high correlation between confounders hindered the sensible estimation of their cumulative effect.

### Impact of potential confounders on trait correlation in couples

Next, we assessed the impact of potential confounders on trait correlation in couples by calculating the ratio of correlation due to confounding over the raw phenotypic correlation among couples averaged across all traits tested (Table 1). While geographical location (using place of birth North/East coordinates) was found to have a negligible impact on phenotypic correlations (mean confounding ratio: 1%), household income (mean confounding ratio: 29.8%), age completed full time education (mean confounding ratio: mean confounding ratio: 11.6%), and physical activity levels (measured using the variable “leisure/social activities: sport club or gym”; mean confounding ratio: 17.1%)) had an important confounding impact on raw phenotypic correlation among couples (Figure 5).

**Figure 5:**
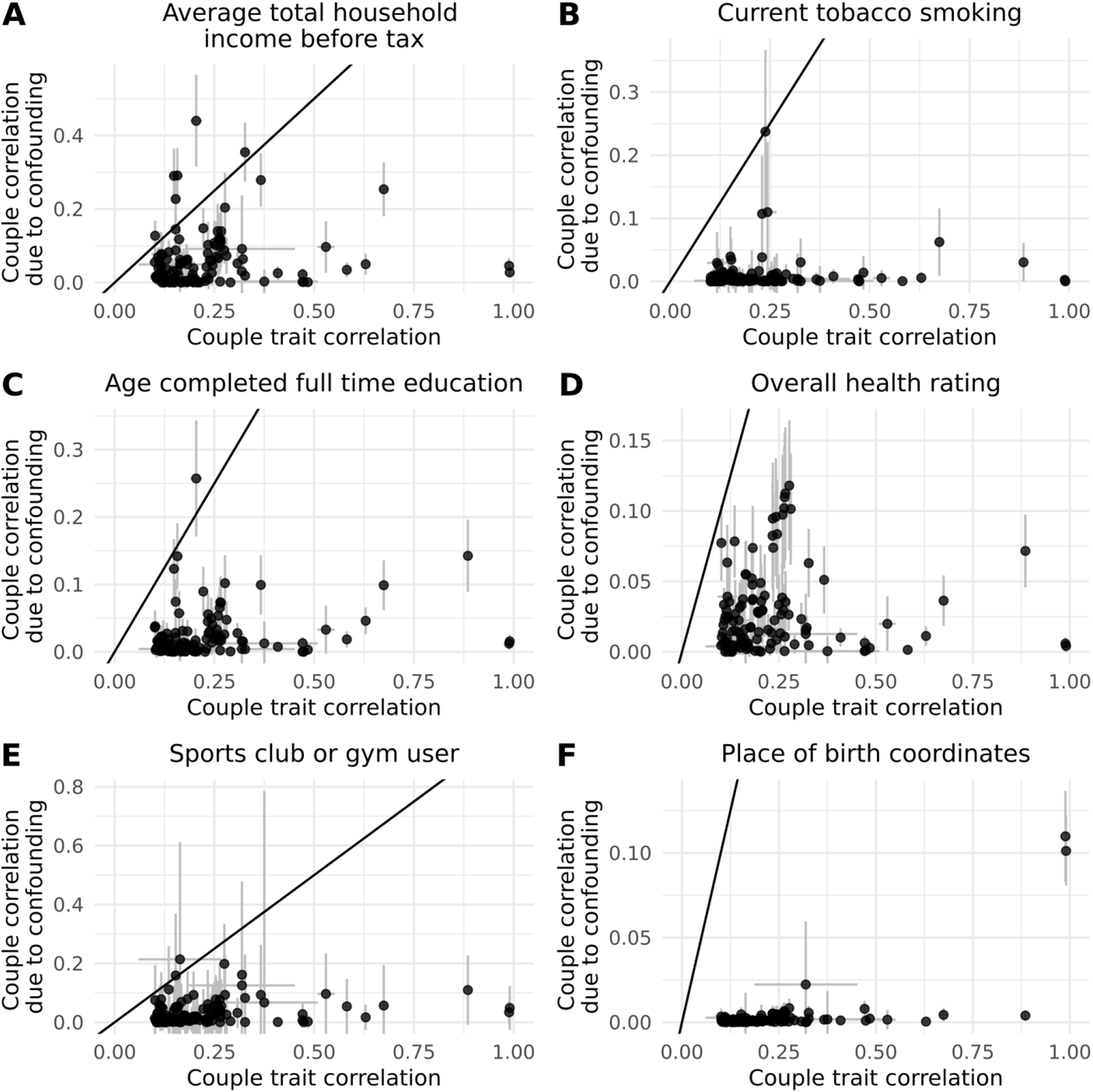
Global confounding impact of select traits on couple phenotypic correlation. Figures display scatter plots of couple correlation due to confounding versus the phenotypic trait correlation among couples for selected potential confounder traits (Z); error bars represent the 95% CI. For each trait in the pipeline, we tested the contribution of four confounder traits (average household income, current tobacco smoking, overall health rating, age completed full-time education, and sports club or gym user, and place of birth co-ordinates, panels **A, B, C, D, E** and **F**, respectively) could impact the phenotypic couple correlation. The couple correlation due to confounding for each trait X was calculated for each confounder Y as C = α_y→x_^2^ × α_y_i_→y_p__. In the case of birthplace coordinates, C-values were summed across the two (independent) North and East coordinates. The identity line is shown in black.

**Table 1:** Summary of the global confounding impact of four selected traits on phenotypic couple correlations. The confounding ratio corresponds to the ratio of correlation due to confounding over phenotypic correlation in couples (i.e. C/cor(X_i_, X_p_), where C = α_y→x_^2^ × α_y_i_→y_p__).

| Confounder trait (Y) | Mean confounding ratio | Median confounding ratio | sd(confounding ratio) |
| --- | --- | --- | --- |
| Current tobacco smoking | 0.047784 | 0.013493 | 0.115313 |
| Overall health rating | 0.141006 | 0.096063 | 0.14611 |
| Leisure/social activities: Sports club or gym | 0.171141 | 0.097595 | 0.215961 |
| Average total household income before tax | 0.298122 | 0.183867 | 0.38749 |
| Age completed full time education | 0.116337 | 0.062482 | 0.177151 |
| Birth place coordinates (North and East) | 0.010319 | 0.005854 | 0.015892 |

### Identification of underlying mechanisms for cross-trait assortment

We sought to identify the mechanisms underlying AM by comparing three estimated paths from a phenotype in the index case (*X_i_*) to another phenotype in its partner (*Y_p_*) as illustrated in Figure 1C. The total causal effect between *X_i_* and *Y_p_* (denoted by *ω*) can be split up into three components: (i) assortative mating through *X* (i.e. *X_i_* → *X_p_*) followed by a causal effect between *X* and *Y* in the partner (i.e. *X_p_* → *Y_p_*), their product being denoted by *γ;* (ii) causal effect between *X* and *Y* in the index individual (i.e. *X_i_* → *Y_i_*), followed by assortative mating through *Y* (i.e. *Y_i_* → *Y_p_*), their product being denoted by *ρ*; (iii) any remaining effect of *X_i_* on *Y_p_*. We computed within-couple cross-trait causal effect estimates *X_i_* → *X_p_* (i.e. 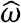) for all combinations of trait pairs (*X, Y*). Of these, we identified 1327 significant MR effects 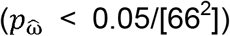 among couples, which were reduced to 1088 pairs after removing pairs with phenotypic correlation > 0.8 (a summary of a set of pruned traits can be found in Supplementary Table 6). Several relationships were almost completely dominated by *ρ* (assortative mating through the outcome), and others dominated byy (assortative mating through the exposure). Specifically, we found 326 relationships which were significantly different between and *γ*, of which 89 (27.3%) showed larger effects through and the other 237 (72.7%) showed larger effects through *γ*. For instance, we found causal relationships between partners for leg fat percentage and time spent watching television; BMI and overall health rating; all dominated by *ρ*. On the other hand, we found some causal relationships between partners which were primarily dominated by *γ* (assortative mating through the exposure), including: comparative height at age 10 (i.e. “When you were 10 years old, compared to average would you describe yourself as: shorter, taller, average”) and forced vital capacity; and standing height on hand grip strength. Finally, we found other pairs where neither *ρ* nor *γ* captured the relationship (i.e. 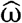 was significantly larger than both estimates), including BMI effect on partner’s systolic/diastolic blood pressure.

Finally, we estimated the contribution of the first two components (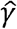 and 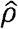) contributing to these significant cross-trait effects, and compared their contribution to the total effect using standard linear regression (Figure 6, Table 2). Paired t-test comparing 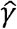 and 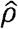 effect estimates revealed that 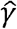 (assortative mating through *X*) is stronger (*p* = 1.1 × 10^5^) in general compared to 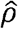 (assortative mating through *Y*). When we summed up the effects of 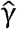 and 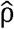, we found that the sum was significantly larger than 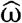. However, these two effects seemed to be correlated, carrying potentially shared signals. Hence, we first residualized 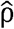 for the effects of 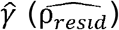, to ensure independence between the two estimates, and then added 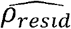 to 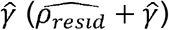. We found no significant difference between 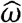 and the sum of 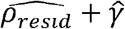 in this analysis and with data points in general, falling near the identity line suggesting that 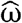 was capturing the paths given by 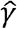 and 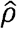. Indeed, linear regression results revealed that 76% of the total effect 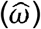 can be explained by the two paths 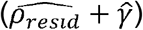 and that the 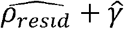 is on average very close to the total effect.

**Figure 6:**
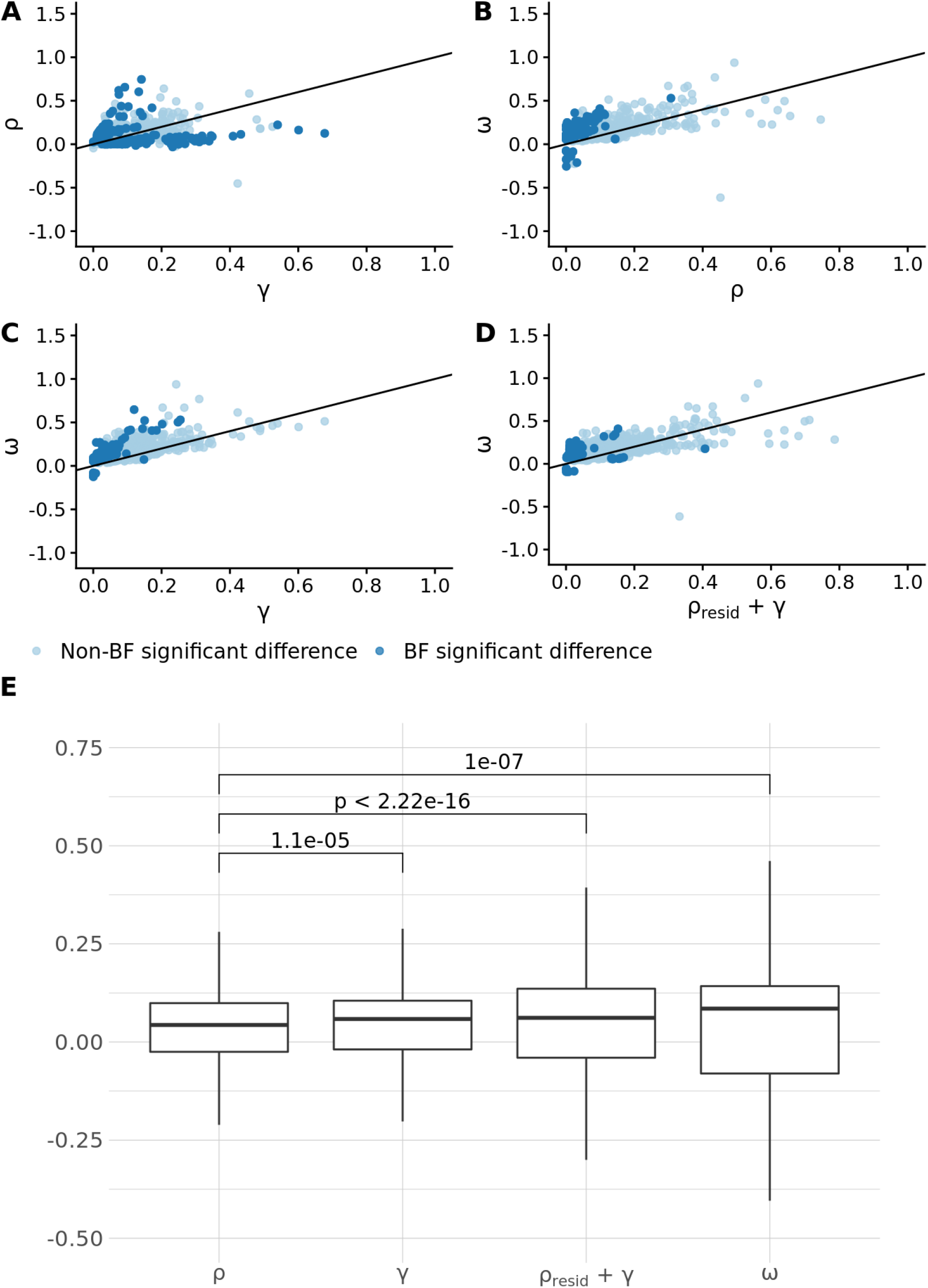
Comparison of causal paths between two traits within couples. Panels **A** through **D** show the regression between the various paths from the index case (*X_i_*) to another phenotype in its partner *Y_p_*) for the 1088 trait pairs with significant MR-effects among couples 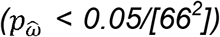 and had correlation < 0.8. To calculate 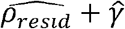, we residualized 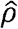 for the effects of 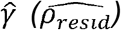, to ensure complete independence between the estimates, and then added 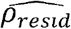 to 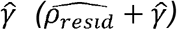. Panel **E** displays a boxplot comparing the coefficients of the estimates among the trait pairs, after removing 19 trait-pairs where the sign did not match between any combination of the four coefficients.

**Table 2:** Summary of the linear regression results given by the model *V* ~ *ω* + 0.

| Variable (V) | $\hat{\beta}$ | SE | p-value | R<sup>2</sup> |
| --- | --- | --- | --- | --- |
| $\hat{\rho}$ | 0.593941 | 0.013169 | 3.2E-251 | 0.651731 |
| $\hat{\gamma}$ | 0.617575 | 0.010114 | < 5E-300 | 0.774278 |
| $\widehat{\rho_{resid}} + \hat{\gamma}$ | 0.769167 | 0.013218 | < 5E-300 | 0.75701 |

## Discussion

In this report, we sought to investigate causal relationships among couples within the UKBB using MR. We analysed 118 traits, representing a wide range of anthropometric-, behavioural-, and disease-related traits. Among the 118 phenotypes tested, we found widespread evidence of causal effects among partners. In particular, we identified 64 same-trait causal effects within partners (out of 118 traits), and no evidence of heterogeneity among same-trait couple MR estimates (*α*_*x_i_*→*x_p_*_). This suggests that associations between the index genotype and partner’s phenotype are primarily acting indirectly through the causal relationship between the traits, rather than the presence of a direct effect for index genotype to the partner’s phenotype. If we assume that genetic effects to partner traits can only happen via first altering a trait of the index case, pleiotropic instruments would only emerge from indirect genetic effects (through another trait), which could be tested and excluded via phenome-wide association studies.

Our results point to AM being stronger among females compared to males which is consistent with the notion first put forth by Darwin, that females are, in general, choosers, while males are courters^31^. Furthermore, our results suggest that fitness and anthropometric measures are important initially (at the time of mate-choice), but their correlation decreases with time, i.e. the longer people stay together the less important it becomes to stay similar in those aspects. On the other hand, we found that smoking cessation and medication use (aspirin, specifically) become more concordant among couples as age increases. As age and time-spent-together are highly correlated variables, it is difficult to distinguish whether this is an effect of convergence or suggestive of an age-dependent mate-choice. We did not identify any significant trends of causal MR effects on time-spent-together or by age. While this could be due to limitations such as statistical power, this is consistent with previous reports which suggest that initial mate choice is a more dominant factor in contributing to phenotypic similarity compared to convergence^7,32–34^.

When investigating the impact of common confounders on our entire panel of phenotypes (i.e. fixing a confounder and assessing its widespread impact on all singletrait aM), we found that household income, age completed education, participant of a sport club or gym are important confounders, explaining on average 29.8, 11.6 and 17.1% of the phenotypic couple correlations among traits tested, respectively. These results also suggest that phenotypic correlations in couples are significantly confounded and point to a relatively few key traits which are driving AM observations. These confounder traits are strongly intertwined and hence correlated, therefore elucidating the key driver is not feasible with the data at hand.

Our findings investigating cross-trait assortment suggest that causal effects from *X_i_* to *Y_p_* are primarily driven by assortative mating through *X* (i.e. *X_i_* → *X_p_*) followed by a causal effect within the partner from *X* to *Y* (i.e. *X_p_* → *Y_p_*). In contrast, a less likely path would be the inverse, whereby the presence of a causal effect from *X* to *Y* in an index case is then followed by *Y* being passed directly from index to partner. These results were expected, as it is more reasonable for couples to influence each other at the exposure level rather than the outcome level, especially since often outcome traits (such as diseases) appear much later than mate-choice.

We found 1088 significant cross-trait causal effects within couples(*ω*), which can be summarized by three categories: (1) driven by assortment on the exposure (*ω* = *γ*) (2) driven by assortment on the outcome (*ω* = *ρ*), and (3) not explained by either (i.e. *ω* being greater than both *ρ* and *γ*). Of note, there were fewer cases in category three, where the causal effect from *X_i_* to *Y_p_* was not captured by *γ* or *ρ*, suggestive of either a direct effect *X_i_* to *Y_p_* or the presence of a confounder variable. An example from the first category, involves a positive causal effect of time spent watching television on BMI driven by the fact that partners causally influence each other with respect to time spent watching television, which in turn has an impact on BMI at the individual level. On the other hand, an example of the second category includes a positive causal relationship from height to education, with a stronger path through *ρ*, representing a path whereby height (a proxy for “dynastic” wealth) increases educational attainment (found previously^35^) within a single individual, and AM subsequently occurs via education level. Finally, as an example for category three, we found a negative causal effect never having smoked on leucocyte count within partners, such that leucocyte count was higher among individuals with partners who smoked. While we also identified a significant effect through *γ* (AM through smoking), the effect was much stronger through ω. These findings suggest that there could be a direct effect from index partner by way of second-hand smoke. These results are consistent with previous work showing higher WBC count in smokers^36^, which might already be achieved by second-hand smoking.

This study has limitations which should be considered. First, to increase statistical power and robustness, we focused on traits available in the UKBB with significant correlation amongst couples and more than 5 valid IVs. As a result, anthropometric traits constituted a larger proportion of our traits under study and represent a large percentage of our significant findings. Other phenotypes, such as behavioural and lifestyle traits, were included but had less statistical power due to lower couple correlation and less IVs. Second, with the current data, we were not able to find strong evidence for couple convergence over time. We did make use of both age and time-together data (proxied by time at the same address) to help shed light on this question, and were able to show that certain traits indeed appear to converge as a couple spends more time together while other traits appear to be more important in the selection process (i.e. true assortative mating). However, to properly assess the question, longitudinal data including measures before couples were together would be best suited to disentangle the complex relationship between assortative mating and convergence. Also, while assortative mating through the exposure (y) and the outcome (*ρ*) represent independent paths from *X_i_* to *Y_p_*, our results suggest that the computed effects using MR estimates are not perfectly independent. This could potentially be due to overlap in genetic instruments, bidirectional causal effect between them or the fact that both estimates depend on the causal effect from *X* to *Y*. To the best of our ability, we tried to mitigate this bias by (i) by using a MVMR approach to remove effects of *X* on *Y* in the calculation of *ρ*, and (ii) first residualising *γ* for effects of *ρ* to ensure independence prior to summation of the effects. Finally, we were limited to the available traits and white British samples in the UKBB. AM is highly population-specific; hence our findings are not necessarily generalisable to other populations.

In summary, we have surveyed a large number of complex traits with significant couple correlation in the UK Biobank and explored to which extent the observed couple similarity is due to couple convergence or confounding. We demonstrated that cross-trait assortment can largely be explained by single-trait assortments between either trait and substantial causal effects between these traits. Our findings provide insights into possible mechanisms underlying observed AM patterns at an unprecedented scale and resolution.

## References

1. Silventoinen, K., Kaprio, J., Lahelma, E., Viken, R. J. & Rose, R. J. Assortative mating by body height and BMI: Finnish twins and their spouses. Am. J. Hum. Biol. 15, 620–627 (2003).

2. Maes, H. H., Neale, M. C. & Eaves, L. J. Genetic and environmental factors in relative body weight and human adiposity. Behav. Genet. 27, 325–51 (1997).

3. Keller, M. C. et al. The Genetic Correlation between Height and IQ: Shared Genes or Assortative Mating? PLoS Genet. 9, (2013).

4. Mare, R. D. Five Decades of Educational Assortative Mating. Am. Sociol. Rev. 56, 15–32 (1991).

5. Agrawal, A. et al. Assortative mating for cigarette smoking and for alcohol consumption in female Australian twins and their spouses. Behav. Genet. 36, 553–566 (2006).

6. Buss, D. M. Marry Someone Who Is Similar To Us in Almost Every Variable. Am. Sci. 73, 47–51 (1985).

7. Watson, D. et al. Match makers and deal breakers: Analyses of assortative mating in newlywed couples. J. Pers. 72, 1029–1068 (2004).

8. Hippisley-Cox, J. Married couples’ risk of same disease: cross sectional study. BMJ 325, 636–636 (2002).

9. Buss, D. M. et al. International Preferences in Selecting Mates. J. Cross. Cult. Psychol. 21, 5–47 (1990).

10. Buss, D. M. & Barnes, M. Preferences in Human Mate Selection. J. Pers. Soc. Psychol. 50, 559–570 (1986).

11. Anderson, C., Keltner, D. & John, O. P. Emotional Convergence Between People over Time. J. Pers. Soc. Psychol. 84, 1054–1068 (2003).

12. Gonzaga, G. C., Campos, B. & Bradbury, T. Similarity, Convergence, and Relationship Satisfaction in Dating and Married Couples. J. Pers. Soc. Psychol. 93, 34–48 (2007).

13. Humbad, M. N., Donnellan, M. B., Iacono, W. G., McGue, M. & Burt, S. A. Is spousal similarity for personality a matter of convergence or selection? Pers. Individ. Dif. 49, 827–830 (2010).

14. Risch, N. et al. Ancestry-related assortative mating in Latino populations. Genome Biol. 10, (2009).

15. Sebro, R., Peloso, G. M., Dupuis, J. & Risch, N. J. Structured mating: Patterns and implications. PLoS Genet. 13, 1–22 (2017).

16. Abdellaoui, A. et al. Genetic correlates of social stratification in Great Britain. Nat. Hum. Behav. 3, 1332–1342 (2019).

17. Rawlik, K., Canela-Xandri, O. & Tenesa, A. Indirect assortative mating for human disease and longevity. Heredity (Edinb). 123, 106–116 (2019).

18. Robinson, M. R. et al. Genetic evidence of assortative mating in humans. Nat. Hum. Behav. 1, 0016 (2017).

19. Xia, C., Canela-Xandri, O., Rawlik, K. & Tenesa, A. Evidence of horizontal indirect genetic effects in humans. Nat. Hum. Behav. 05, 399–406 (2020).

20. Yengo, L. et al. Imprint of assortative mating on the human genome. Nat. Hum. Behav. 2, 948–954 (2018).

21. Border, R. et al. Assortative mating biases marker-based heritability estimators. Nat. Commun. 13, (2022).

22. Border, R. et al. Cross-trait assortative mating is widespread and inflates genetic correlation estimates. bioRxiv 2022.03.21.485215 (2022) doi:10.1101/2022.03.21.485215.

23. Lawlor, D. A. et al. Mendelian randomization: using genes as instruments for making causal inferences in epidemiology. Stat. Med. 27, 1133–1163 (2008).

24. Howe, L. J. et al. Genetic evidence for assortative mating on alcohol consumption in the UK Biobank. Nat. Commun. 10, (2019).

25. Burgess, S., Butterworth, A. & Thompson, S. G. Mendelian randomization analysis with multiple genetic variants using summarized data. Genet. Epidemiol. 37, 658–665 (2013).

26. Millard, L. A. C., Davies, N. M., Gaunt, T. R., Smith, G. D. & Tilling, K. Software application profile: PHESANT: A tool for performing automated phenome scans in UK Biobank. Int. J. Epidemiol. 47, 29–35 (2018).

27. Pirastu, N. et al. Using genetic variation to disentangle the complex relationship between food intake and health outcomes. bioRxiv 829952 (2020) doi:10.1101/829952.

28. Burgess, S., Small, D. S. & Thompson, S. G. A review of instrumental variable estimators for Mendelian randomization. Stat. Methods Med. Res. 26, 2333–2355 (2017).

29. Gao, X., Starmer, J. & Martin, E. R. A multiple testing correction method for genetic association studies using correlated single nucleotide polymorphisms. Genet. Epidemiol. 32, 361–369 (2008).

30. Hemani, G., Tilling, K. & Davey Smith, G. Orienting the causal relationship between imprecisely measured traits using GWAS summary data. PLOS Genet. 13, e1007081 (2017).

31. Ryan, M. J. Darwin, sexual selection, and the brain. Proc. Natl. Acad. Sci. U. S. A. 118, 1–8 (2021).

32. Mascie-Taylor, C. G. N. Spouse similarity for IQ and personality and convergence. Behav. Genet. 19, 223–227 (1989).

33. Caspi, A., Herbener, E. S. & Ozer, D. J. Shared experiences and the similarity of personalities: a longitudinal study of married couples. J. Pers. Soc. Psychol. 62, 281–91 (1992).

34. Yengo, L. et al. No Evidence for Social Genetic Effects or Genetic Similarity Among Friends Beyond that Due to Population Stratification: A Reappraisal of Domingue et al (2018). Behav. Genet. 50, 67–71 (2019).

35. Tyrrell, J. et al. Height, body mass index, and socioeconomic status: Mendelian randomisation study in UK Biobank. BMJ 352, (2016).

36. Pedersen, K. M. et al. Smoking and Increased White and Red Blood Cells: A Mendelian Randomization Approach in the Copenhagen General Population Study. Arterioscler. Thromb. Vasc. Biol. 39, 965–977 (2019).

