## Supplementary Information for "The contribution of mate-choice, couple convergence and confounding to assortative mating"

### Supplementary Figures

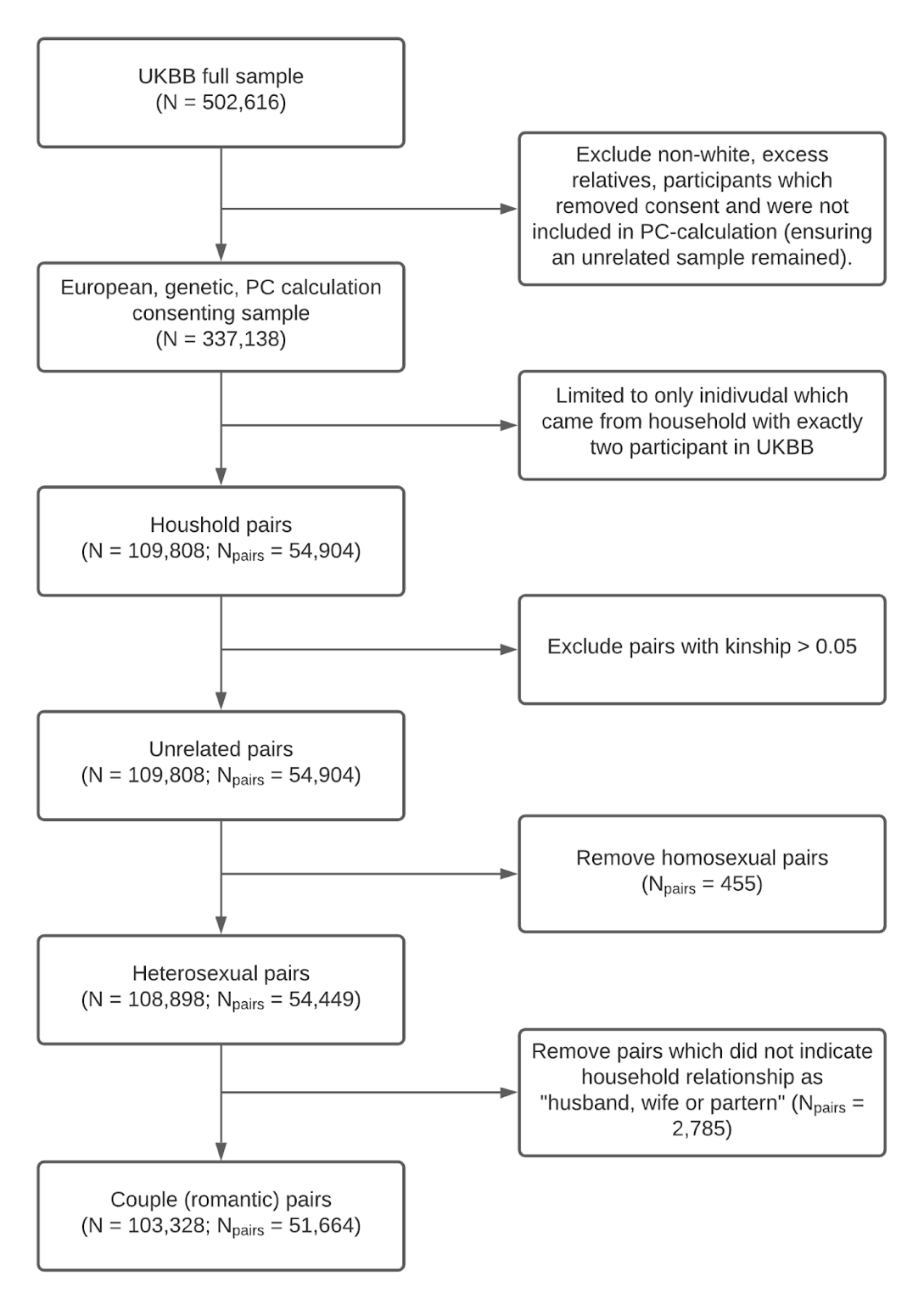

**Supplementary Figure 1:** **Couple selection summary.** *Flow chart shows a summary of the couple determination and selection in the UKBB.*

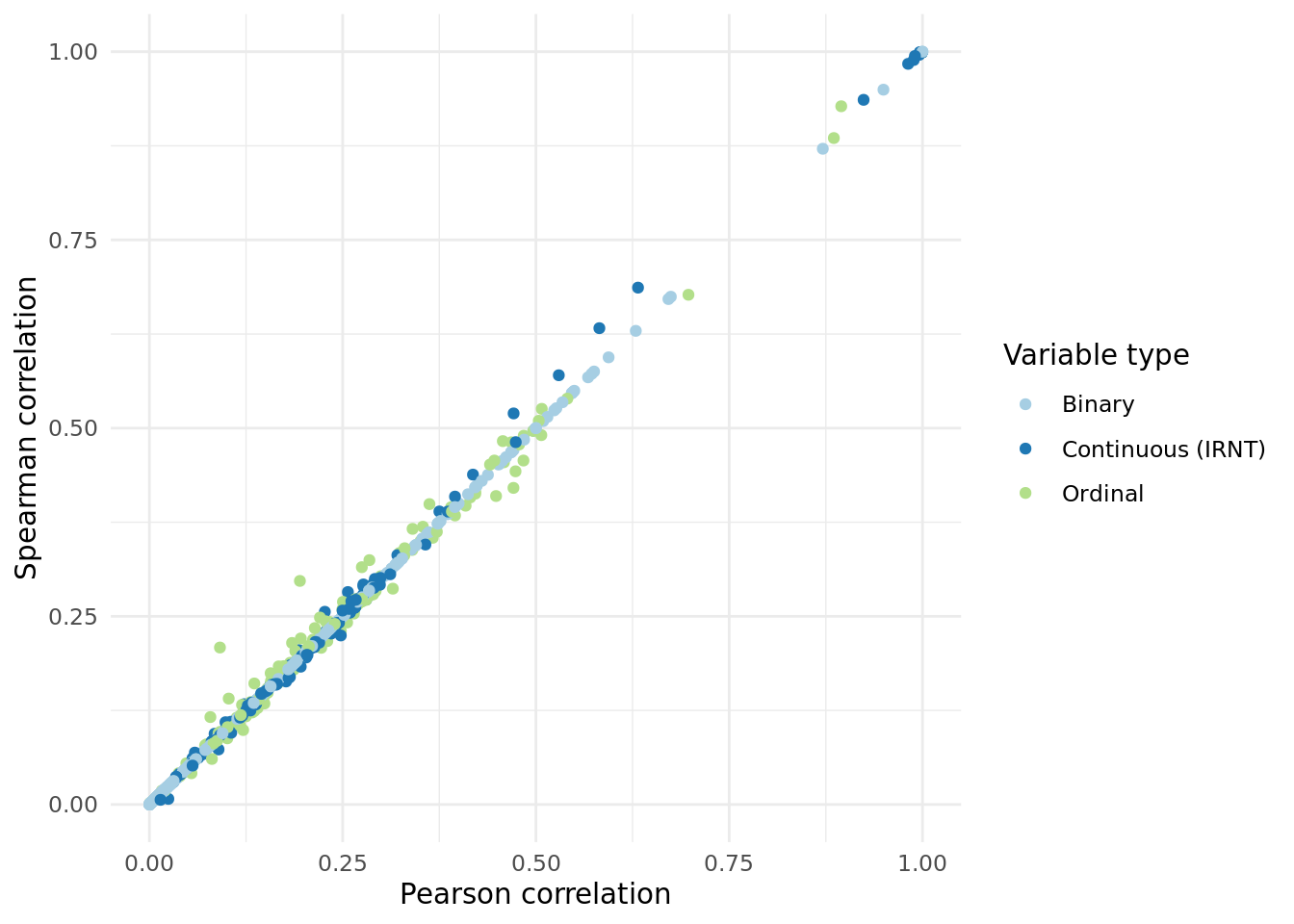

**Supplementary Figure 2:** **Spearman versus Pearson correlation in the UKBB.** *Scatter plot shows Spearman correlation versus Pearson correlation for each trait considered (n = 1278), color coded according to variable type. Consistent correlation estimates were obtained regardless of variable type.*

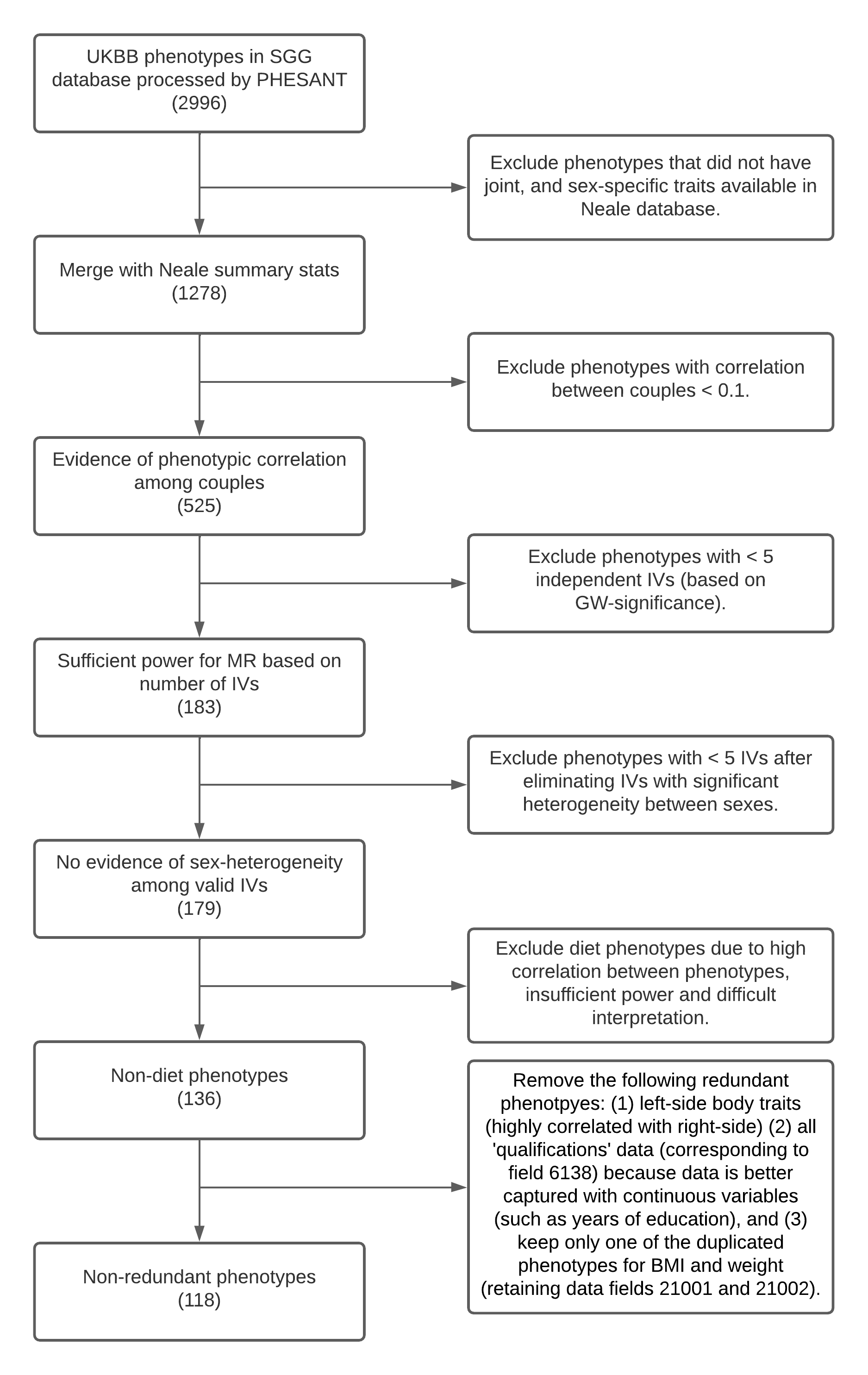

**Supplementary Figure 3:** **Phenotype selection summary.** *Flow chart shows a summary of the phenotype selection included in the pipeline, resulting in 118 phenotypes for analysis. SGG refers to the Statistical Genetics Group, data was used from the corresponding internal UKBB database under application number #16389.*

### Supplementary Table Descriptions

**Supplementary Table 1:** Summary of the Bonferroni-significant single-trait MR results within couples (*p* < 0.05/66).

**Supplementary Table 2:** Summary of the nominally-significant MR sex differences (Z-test, *p* < 0.05) among the 64 traits that showed a significant overall MR-effect (in both sexes combined).

**Supplementary Table 3:** Summary of the Bonferroni-significant traits which showed a significant linear trend of Pearson phenotype correlations versus bins by age or time-spent-together (*p* < 0.05**/**66).

**Supplementary Table 4:** Summary of the Bonferroni-significant traits which showed significant differences between MR-estimates and phenotypic correlations amongst couples (Z-test, p < 0.05/66). Of the significant differences, 40 had larger phenotypic correlations compared MR-estimates, and 3 traits showed larger MR-estimates compared to phenotypic correlation.

**Supplementary Table 5:** Summary of the identified potential confounders for the traits which showed phenotypic correlations significantly larger than MR-estimates. Of the 40 traits investigated, we were able to identify at least 1 potential confounder for all but one.

**Supplementary Table 6:** Summary of the significant two-trait MR associations ($p_{\hat{\omega}}$ < 0.05/[66^2^]). Results are pruned such that pair A-B and C-D and if max(corr(A,C)*corr(B,D),corr(A,D)*corr(B,C)) > 0.8 then one pair is dropped, prioritized by lower $p_{\hat{\omega}}$.
